# Converting endogenous genes of the malaria mosquito into simple non-autonomous gene drives for population replacement

**DOI:** 10.1101/2020.05.09.086157

**Authors:** Astrid Hoermann, Sofia Tapanelli, Paolo Capriotti, Ellen K. G. Masters, Tibebu Habtewold, George K. Christophides, Nikolai Windbichler

**Affiliations:** Department of Life Sciences, Imperial College London, London, United Kingdom

**Keywords:** population replacement, malaria, CRISPR/Cas9, gene drives

## Abstract

Gene drives for mosquito population replacement are promising tools for malaria control. However, there is currently no clear pathway for safely testing such tools in endemic countries. The lack of well-characterized promoters for infection-relevant tissues and regulatory hurdles are further obstacles for their design and use. Here we explore how minimal genetic modifications of endogenous mosquito genes can convert them directly into non-autonomous gene drives without disrupting their expression. We co-opted the native regulatory sequences of three midgut-specific loci of the malaria vector *Anopheles gambiae* to host a prototypical antimalarial molecule and guide-RNAs encoded within artificial introns, that support efficient gene drive. We assess the propensity of these modifications to interfere with the development of *Plasmodium falciparum* and their effect on fitness. Because of their inherent simplicity and passive mode of drive such traits could form part of an accepted testing pathway of gene drives for malaria eradication.

## Introduction

After more than a decade of sustained success in the fight against malaria, data from 2015 onwards suggests that no significant progress in reducing global malaria cases has been achieved (1). The rise of insecticide resistance in mosquito vectors and drug resistance in parasites highlights the urgent need to develop new tools if malaria eradication is to remain a viable goal.

Synthetic gene drives spreading by super-Mendelian inheritance within vector populations have been proposed as an area-wide genetic strategy for the control of malaria (2). They mimic the mechanism of proliferation of a class of naturally occurring selfish genes called homing endonucleases and could be deployed to modify the genetic makeup of disease vector populations (3, 4). Proof-of-principle laboratory experiments of CRISPR/Cas9-based gene drives for population suppression, which aims at the elimination of the target population (5, 6), as well as population replacement (7, 8), which aims to spread an anti-malarial trait within a vector population, suggest that these strategies could be deployed to reduce malaria transmission in the field.

The success of population replacement in particular hinges on the availability of molecules that can efficiently block *Plasmodium* development within the mosquito vector as well as ways to express such elements. Few pre-characterized promoter elements, driving tissue-specific expression in infection-relevant tissues currently exist in malaria vectors. Examples include the regulatory elements of the zinc carboxypeptidase A1 (CP), peritrophin1 (Aper1) or the vitellogenin (Vg) genes that have been reported to drive transgene expression (9-12). A variety of endogenous and exogenous effector molecules have been expressed in transgenic mosquitoes to interfere with the development of the parasite including anti-malarial regulators such as Cecropin A (13), Fibrinogen domain-containing immunolectin 9 (FBN9) (14), TEP1 (15), the transcription factor Rel2 (16), as well as the protein kinase Akt (17, 18) and the Phosphatase and Tensin homolog (PTEN) (19) involved in insulin-signalling. Synthetic peptides Vida3 (10) and SM1, engineered as a quadruplet (20), the bee venom phospholipases A2 (PLA2) (11, 21), as well as single-chain antibodies (scFvs, m1C3, m4B7, m2A10) targeting the ookinete protein Chitinase 1 and the Circumsporozoite protein (CSP) (22, 23) have been used. Recently, microRNA sponges have been suggested as modulators of mosquito immunity (24). The performance of these effector molecules has been evaluated using laboratory strains of the human parasite *Plasmodium falciparum* or the rodent parasite *Plasmodium berghei*, but the efficacy against circulating polymorphic *P. falciparum* strains of human malaria parasites is currently unknown.

A further consideration is the population dynamics and persistence of transmission-blocking gene drives (25, 26) which would suggest a trade-of between the targeting of a conserved sequence with a resulting fitness cost versus the targeting of a neutral genomic site that shows poor conservation and engenders the evolution of resistance to the drive. A number of ways have been suggested to work around this, for example the linking of rescue-copies of essential genes to drive elements (27). Cleave and rescue elements are a related approach that does not rely on homing (28, 29). However, not only could such approaches increase molecular complexity of the constructs (and likely also their regulatory approval) but it could also complicate the deployment of multiple waves or combinations of drive and effector within a population.

We have recently modelled an alternative approach to achieve population replacement, that was specifically designed to address these questions (30). The approach we have termed integral gene drive (IGD) imitates the way naturally occurring homing endonuclease genes propagate i.e. by copying themselves into highly conserved sites within genes. They do not disrupt their target genes due to their association with introns or inteins (31, 32). We analysed the theoretical behaviour of IGDs featuring the complete molecular separation of drive and effector functions into minimal modifications of endogenous host genes at two or more different genomic loci (30). This approach takes advantage of the promoter and also of its surrounding regulatory regions of the modified endogenous locus. To accommodate the guide-module required for subsequent gene drive and a marker-module required for monitoring transgenesis, we suggested to insert an intron into the effector gene.

In the present study we sought to scope the feasibility of this approach in the African malaria mosquito *A. gambiae*. Our aim was to generate minimal genetic modifications of mosquito genes *in-situ*, which would turn particular alleles into non-autonomous gene drives whilst also expressing a putative antimalarial effector molecule. We sought to assess the efficacy of gene drive, the ability to express anti-*Plasmodium* molecules by linkage to genes active in infection-relevant tissues and how these modifications would affect the expression of such host genes as well as overall mosquito fitness. In order to establish this experimental proof-of-principle for integral gene drive, we have chosen as a prototypical effector molecule a known anti-microbial peptide (AMP), Scorpine, as it had previously been shown to have a transmission blocking effect on *P.falciparum* in the context of paratransgenesis with several different microorganisms (33-36). For the same reason we focused here on female mid-gut specific genes since they allow targeting the *Plasmodium falciparum* parasite during the ookinete to oocyst transition stage (37).

## Results

### Direct modification of three *A. gambiae* midgut loci

We chose CP (AGAP009593) and Aper1 (AGAP006795) for experimental validation of the IGD strategy because their promoters had been used previously for conventional transgene overexpression (10, 11). CP is expressed in the guts of pupae and sugar-fed adults and becomes ten-fold upregulated by blood-feeding as well as by feeding on protein-free meals, probably triggered by the gut distention (38). Its mRNA levels peak 3 hours post blood-feeding (pbf), with spatial expression limited to the posterior midgut, and return to normal levels 24 hours pbf. Aper1 is only detected in the adult gut, but not in larvae, pupae or adult carcass, and its mRNA expression profile is independent of age and blood-feeding (39). The protein is stored in secretory vesicles and released into the midgut lumen soon after bloodmeal, where it is cross-linked into the peritrophic matrix via its two tandem domains that bind chitin (40). The peritrophic matrix retracts from the midgut wall 48 hours pbf and is excreted after 72 hours pbf (41). Using the VectorBase expression explorer (42) we identified further genes with significant expression in the female midgut and upregulation after the blood-meal. Amongst them we chose alkaline phosphatase 2 (AP2, AGAP006400), which has a secretion signal and a GPI-anchor and is detected in the detergent resistant membrane (DRM) proteome of non-blood-fed adult *Anopheles gambiae* midguts (43).

We next identified guide RNA (gRNA) target sites around the start and stop codons of these genes for the insertion of the construct sequences (Figure 1A). The gRNAs were chosen based on a compromise between proximity of the cut site to the start or stop codons, activity and off-target scores as well as the lack of common target site polymorphisms in public population genetic datasets of the mosquito (Table S1) and all gRNAs were evaluated *in vitro*.

**Figure 1.**
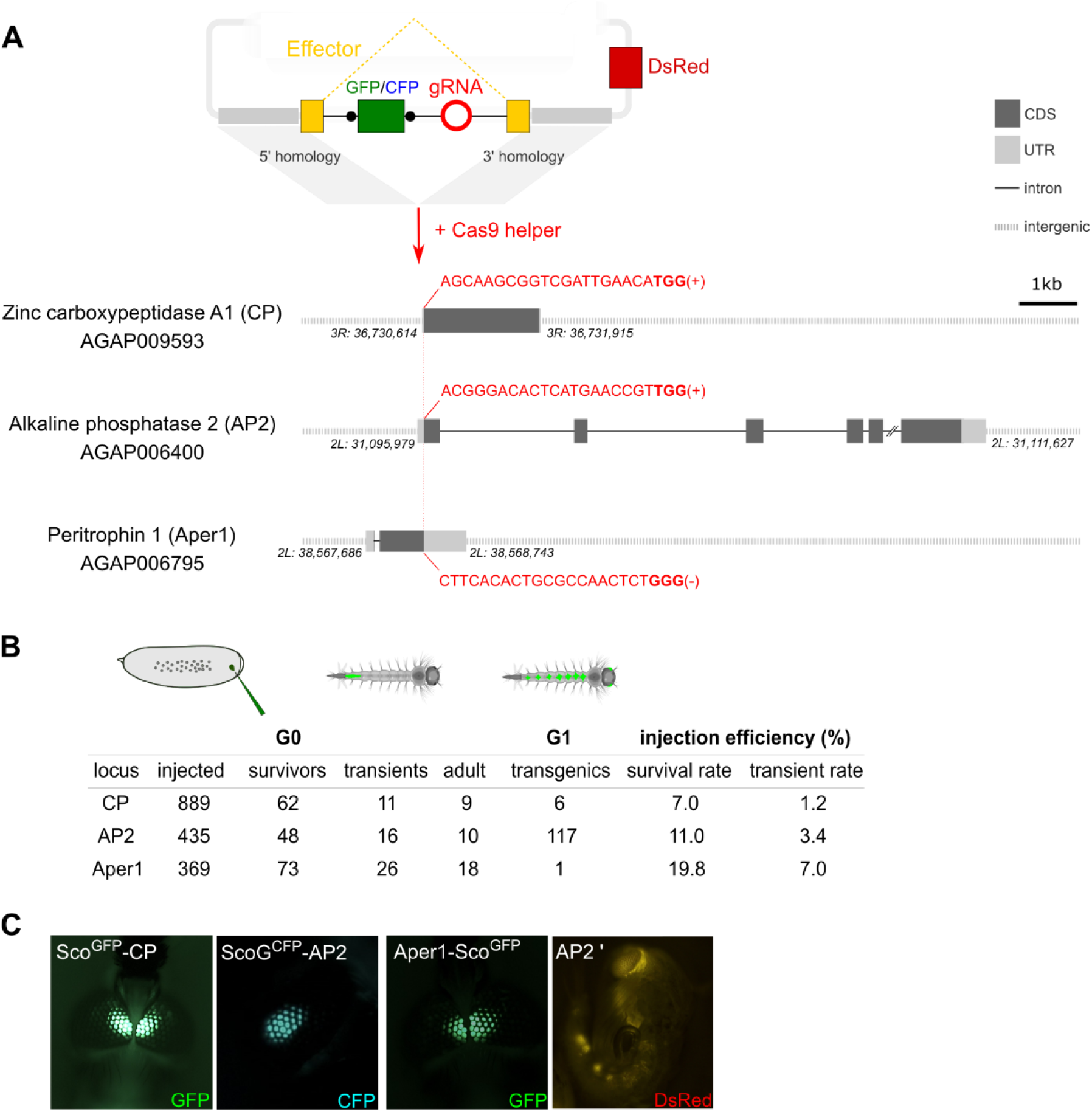
Homology-directed modification of *Anopheles gambiae* midgut loci by CRISPR/Cas9. **(A)** Schematic showing the insertion of the effector construct at the Carboxypeptidase (CP), Alkaline phosphatase 2 (AP2) and Peritrophin 1 (Aper 1) loci. The donor plasmid supplies the effector coding sequence (yellow), which accommodates an artificial intron. The intron harbours a fluorescent marker (either GFP or CFP, green) under the control of the 3xP3 promoter flanked by loxP sites (black dots) and a U6 driven guide-RNA module (red), required for both transgenesis and subsequently for gene drive. The plasmid features regions of homology that drive N- or C-terminal insertion at the start (CP, AP2) or stop codon (Aper1) as well as a 3xP3::DsRed plasmid-backbone marker. The gRNA target sequence (red) including the PAM motif (bold), and the target strand are indicated. **(B)** Summary of embryo microinjections and the identification of transgenic individuals by fluorescent screening. **(C)** Adult transgenic mosquitos with fluorescent expression of GFP or CFP in the eyes under the control of the 3xP3 promoter as well as a pupa showing DsRed fluorescence, indicating plasmid-backbone integration (AP2’).

We generated transformation constructs consisting of homology arms for these 3 loci designed to facilitate integration of the Scorpine coding sequence within each gene. In the case of CP and AP2, the Scorpine coding sequence was inserted at the start codon and linked to the coding sequence of the host gene via the 2A autocleavage-peptide (44). In contrast, we anchored the effector to the peritrophic matrix via a C-terminal fusion to Aper1. In the AP2 strain, Scorpine is expressed as a GFP-fusion with the aim to enhance stability of the protein in the protease enriched gut environment. In each case, the effector coding sequences harboured an artificial intron encoding the gRNA (Figure 1A). We have recently characterized, in S12 cells and transgenic *Drosophila* strains, a number of artificial introns optimized equally for splicing and gRNA expression using an intronic RNA polymerase III promoter (A. Nash et al. 2020, in preparation) and we applied these designs here. Each artificial intron harbours a fluorescent marker driven by the 3xP3 promoter, flanked by loxP-sites, and the corresponding guide RNA under the control of the ubiquitous and constitutive *Anopheles gambiae* U6 promoter (Figure 1A).

For transgenesis, we co-injected a helper plasmid carrying the Cas9 coding sequence under the control of the vasa promoter (6) and the results of these experiments are summarized in Figure 1B and C. We established the three transgenic strains Sco^GFP^-CP, ScoG^CFP^-AP2 and Aper1-Sco^GFP^ in the latter case from a single G1 founder (Figure 1B). Integration of the vector backbone distinguished via the additional DsRed marker was only observed for the AP2 locus in several G1 individuals that were discarded (Figure 1C). Sequencing of all individuals within a G1 founder-cage for each strain confirmed that insertion into the genomic locus was precise at all 3 loci, and no aberrant integration events could be detected.

### Establishing minimal genetic modifications

We predicted that efficient splicing of the artificial introns would occur only in the absence of the 3xP3-GFP or CFP fluorescent transformation marker genes, and we thus expected that the integrations we had established would interfere with the function of the mosquito host genes. However, following sib-mating we found that for all 3 modified loci, homozygous individuals were viable and fertile and showed no striking fitness defects during rearing and maintenance. In order to establish minimal genetic modifications, i.e. to remove the fluorescent transformation markers flanked by loxP sites, we next crossed all 3 transgenic strains to a vasa-Cre strain (45). We hereby established the markerless strains Sco-CP, ScoG-AP2 and Aper1-Sco (Figure 2A), all of which were also found to be homozygous viable and fertile. Establishing and tracking markerless modifications in *A. gambiae* is challenging and has not been attempted previously. Specifically, we achieved to establish pure-breeding markerless strains either by genotyping of pupal cases of individuals lacking visible fluorescent markers in the case of Sco-CP (Figure 2B) or in the case of Aper1-Sco by crossing the transhemizygotes to a Cas9 expressing strain to induce homing and preferential inheritance of the markerless allele (Figure 2 – figure supplement 1A). For generating ScoG-AP2, we used the homozygous ScoG^CFP^-AP2 as a dominant balancing marker for tracking inheritance of the unmarked allele (Figure 2 – figure supplement 1B). PCR on genomic DNA (Figure 2C) and subsequent sequencing confirmed precise removal of the marker module, and also suggested successful homing of the construct.

**Figure 2.**
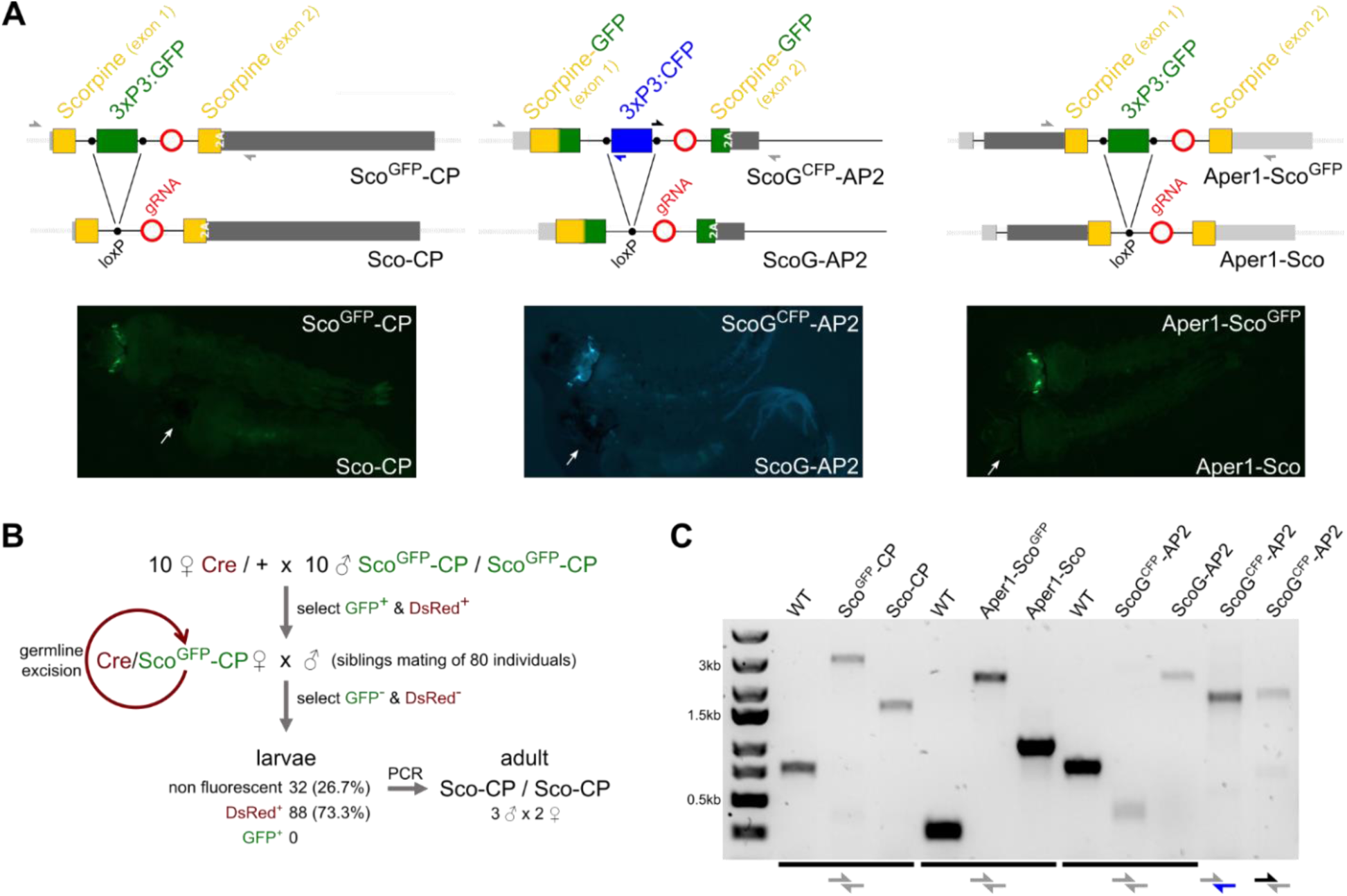
Generation of minimal genetic modifications. **(A)** Schematic showing the inserted transgene constructs within the exon structure of the CP, AP2 and Aper1 loci prior and following the excision of the marker gene by Cre recombinase (top) and the observed changes in green or cyan fluorescence in L3 larvae (bottom). Half arrows indicate primers for the PCRs shown in panel C and white arrows indicate the eyes in the markerless individuals. 2A indicates the F2A self-cleaving peptide signal. **(B)** Crossing scheme used for the establishment of markerless strain Sco-CP by crossing to a Cre recombinase expressing strain. Non-fluorescent adults were allowed to hatch individually, and their pupal cases were used for genotyping. **(C)** PCR genotyping of genomic DNA of homozygous individuals of all strains with primer-pairs (shown in A) spanning the 3 loci. The entire locus could not be amplified in strain ScoG^CFP^-AP2 that contains both GFP and CFP, and hence separate 5’ and 3’ fragments were analysed by PCR. The following figure supplements are available for figure 2. Figure supplement 1. Crossing schemes used for the establishment of markerless strains.

### Expression analysis of integrated effector sequences and their effect on the host genes

For a comparative expression study, we extracted RNA from female midguts 3 hours after blood-feeding of the 3 strains Sco^GFP^-CP, ScoG^CFP^-AP2 and Aper1-Sco^GFP^ and of the corresponding markerless strains Sco-CP, ScoG-AP2 and Aper1-Sco which carry minimal genetic modifications (Figure 3A). RT-PCRs over the splice junctions and subsequent sequencing confirmed that for all 3 markerless strains precise splicing occurs and that the removal of the artificial intron restores the open reading frame required for expression of both, the effector protein and the host gene product (Figure 3B). We further evaluated mRNA expression of the transgenes in the CP and AP2 locus by RT-PCR using primer pairs targeting the N-terminal part of Scorpine preceding the intron or the entire GFP, primers spanning the splice site, as well as primers targeting the host gene coding sequence (Figure 3C, D). Because integration of the transgene sequences is C-terminal in the case of Aper1, we performed qPCR with one primer pair targeting the endogenous Aper1 and another one the region between Aper1 and Scorpine (Figure 3E). Overall, our analysis showed that in all three cases the expression of the downstream (CP or AP2) or upstream (Aper1) host gene is severely reduced by the presence of the 3xP3 fluorescence marker cassette. However, removal of the marker gene restores host gene expression in all 3 strains (Figure 3C, D, E). The transcript for the exogenous effector Scorpine was efficiently detected and amplification over the splice site of the artificial intron was successful in the markerless strains Sco-CP, ScoG-AP2 and Aper1-Sco. We compared Scorpine expression levels between the 3 markerless transgenic strains, before and 3h after blood-feed (Figure 3 – figure supplement 1A). In the case of the integration into AP2, the qPCR signal of Scorpine was at the detection limit in blood-fed guts and low in non-blood-fed guts. For Scorpine within the CP locus, an upregulation after blood-feed was observed in accordance with published data for the CP promoter (38). The highest expression levels were achieved with the integration into Aper1 under both conditions. We also performed mass-spectrometry (LC-MS/MS) on non-blood-fed guts to confirm that protein expression of the co-opted host gene is unaffected by effector integration or splicing of the artificial intron. We detected high-confidence peptides for both genes in the Sco-CP and Aper1-Sco strains (Figure 3 – figure supplement 1B). For Aper1, the peptides map to two out of the three regions from published data, and for Sco-CP 4 out of the 14 published peptides were identified (46).

**Figure 3.**
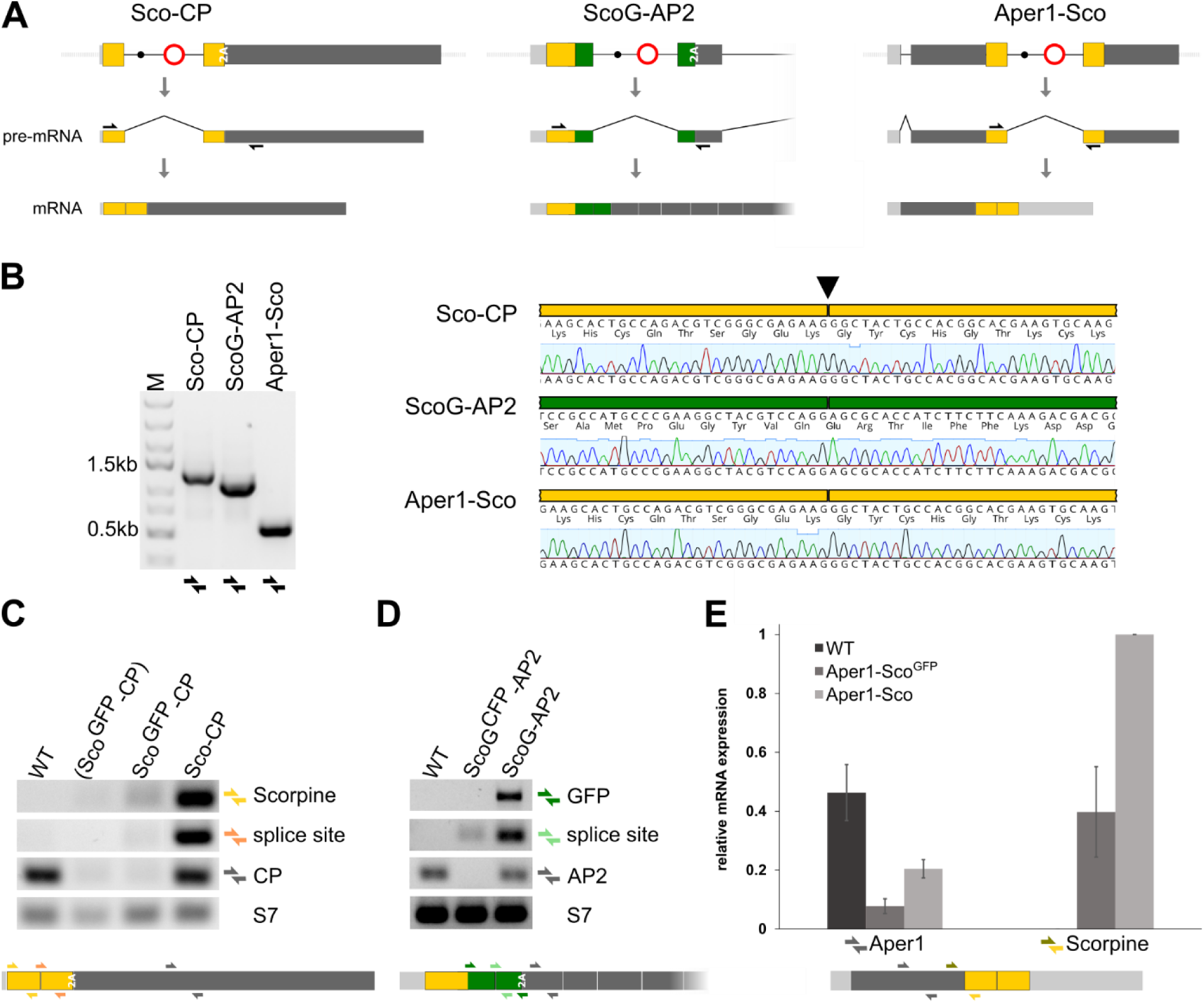
Splicing of the artificial intron and expression analysis. **(A)** Schematic showing mRNA expression of the modified host genes of the 3 markerless strains assuming correct splicing of the artificial intron. Black arrows indicate RT-PCR primers used in B. **(B)** RT-PCR (left) and sequencing of the amplicons (right) indicates precise splicing of the 3 artificial introns. Midguts of homozygous strains Sco-CP, ScoG-AP2 and Aper1-Sco were dissected 3 hours after blood feeding and RNA was extracted for RT-PCR. Analysis of mRNA expression of the modified and unmodified CP **(C)** and AP2 **(D)** loci by RT-PCR. Guts from 10-15 homozygous individuals with or without the marker cassette were compared to the wild type (WT). Pool (Sco^GFP^-CP) was enriched for the transgene but possibly contained hemizygous individuals. Primer pairs are indicated as coloured half arrows in the schematic below. Splice site refers to a primer pair, where one of the primers spans the splice junction. S7 served as house-keeping reference gene. RNA was extracted from midguts 3 hours after the bloodmeal. **(E)** Analysis of mRNA expression of the modified Aper1 locus by qPCR. Guts from 10-15 homozygous individuals with (Aper1-Sco^GFP^) and without (Aper1-Sco) the marker gene were analysed. qPCR with the indicated primer pairs (coloured half arrows) was conducted on cDNA from midguts dissected 3 hours after blood-feed and expression was normalized to the S7 reference gene. Data derive from 3 biological replicates with 3 technical replicates each. The following figure supplements are available for figure 3. Figure supplement 1. Expression levels and mass-spectrometry of co-opted host genes.

### Mosquito fitness analysis

We assessed the effects on fecundity and larval hatching of homozygous strains Sco-CP, ScoG-AP2 and Aper1-Sco and compared them to individuals of the wild type (G3) colony and additionally to the vasa-Cre strain (45), which derives from the *Anopheles gambiae* KIL background (20) (Figure 5A, B). Females of all transgenic strains laid a significantly reduced number of eggs. The Aper1-Sco strain also showed a statistically significant reduction in the hatching rate compared to the G3 wild type (Figure 5B). Because all 3 modified host genes are female and midgut specific and because fitness effects would be expected to be sex-specific we also measured pupal sex ratio of these strains but found no significant deviations from an expected 1:1 sex ratio for any of the transgenics (Figure 5 – figure supplement 1). Because similar effects were observed for all transgenic strains (including the Cre strain) and because of high variability between individual females we concluded that inbreeding effects, rather than effects of the respective genetic modifications, best explain the observed reduction in fertility. Nevertheless, the presence of negative fitness effects of these traits under these or other conditions could not be ruled out by our experiments.

**Figure 4.**
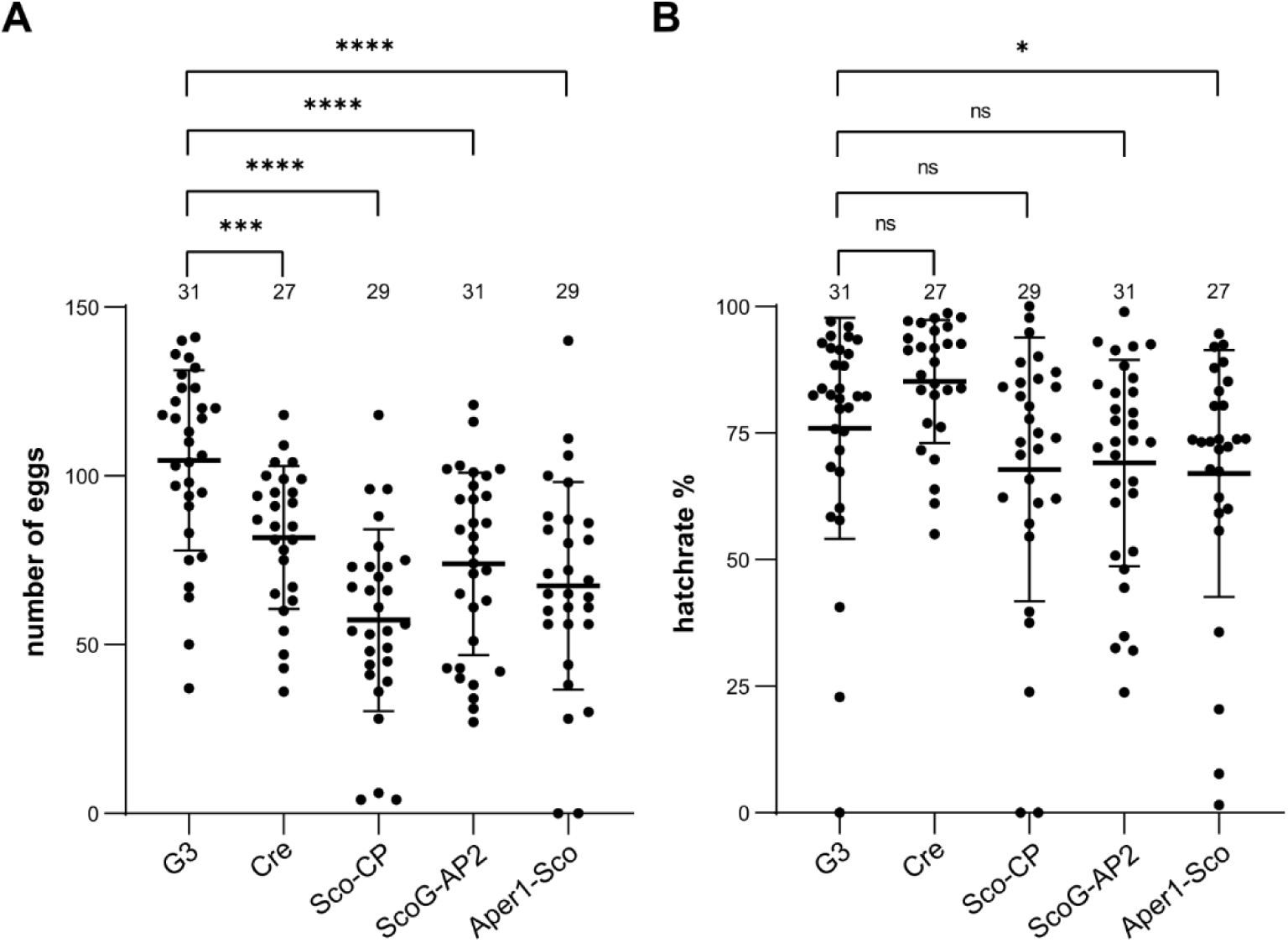
Fecundity and larval hatch rates of the markerless transgenic strains. **(A)** Fecundity of single females of the homozygous markerless strains Sco-CP, ScoG-AP2 and Aper1-Sco compared to the G3 wild type and the vasa-Cre strain (KIL background) used to remove the marker module. Mean and standard deviation are plotted and the total number analyzed from three pooled biological replicates is indicated on top. All datasets have a Gaussian distribution according to the Shapiro-Wilk test. p-values were calculated using the unpaired two-tailed Student’s t-test. **(B)** Larval hatch rates from single females of the markerless strains and the two control strains. None of the five datasets showed a Gaussian distribution according to Shapiro-Wilk, and the p-values were calculated using the Kolmogorov-Smirnov test. * P ≤ 0.05, ** P ≤ 0.01, ***P ≤ 0.001, **** P ≤ 0.0001 and ns not significant. The following figure supplements are available for figure 4. Figure supplement 1: Pupal sex ratio.

**Figure 5.**
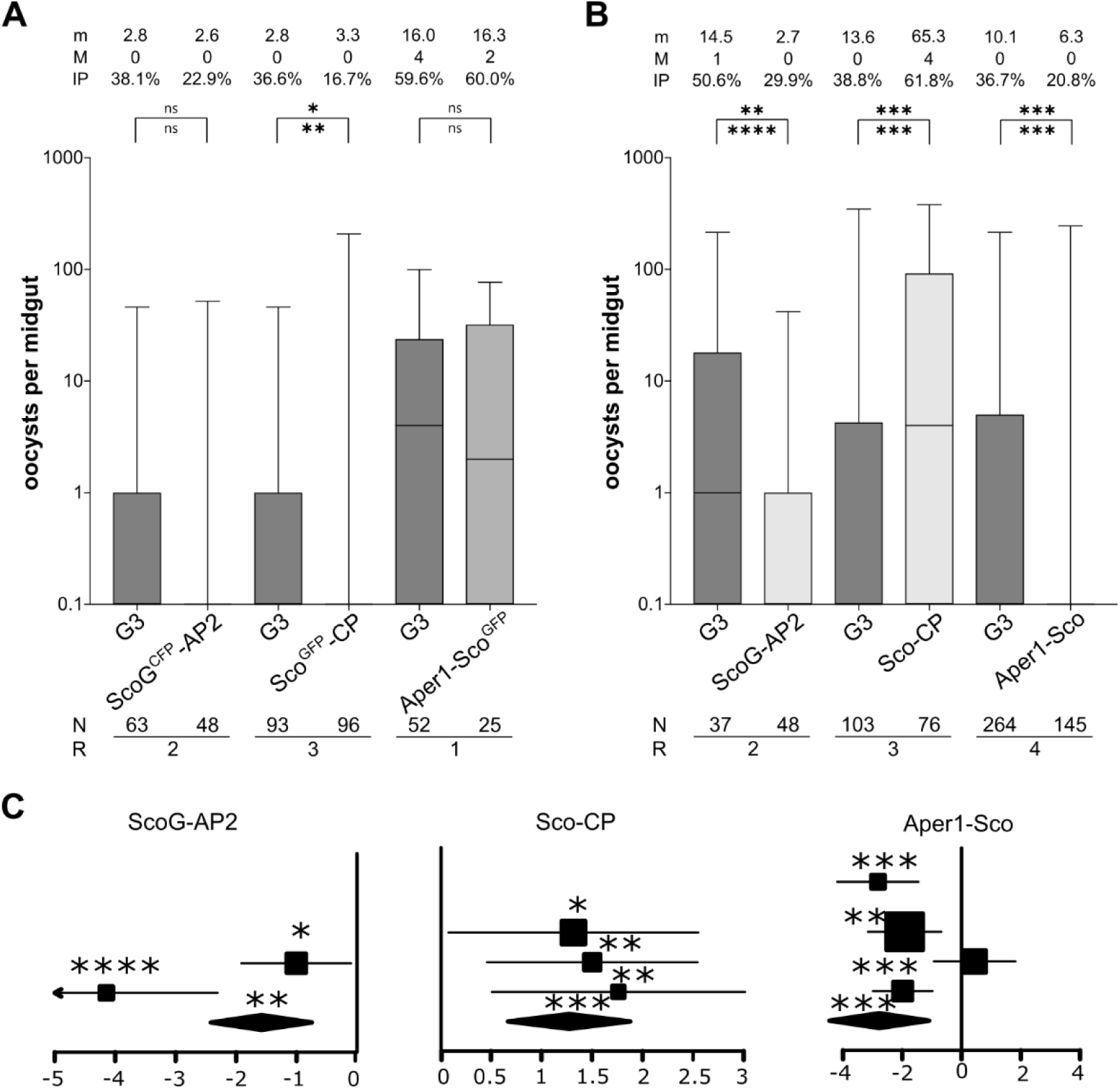
Transmission blocking assay. Standard membrane feeding assay with *P. falciparum* using the homozygous transgenic strains with the marker-module **(A)** and the corresponding markerless strains **(B)**. Infection intensity is measured by the number of oocysts in each gut and the mean (m) and median (M) are shown on top, as well as the infection prevalence (IP). The statistical significance of the infection intensity (stars above the bar) and infection prevalence (below) were calculated with the Mann-Whitney test and the Chi-squared test, respectively. N is the number of mosquitoes analysed and R is the number of replicates performed. **(C)** Analyses of data plotted in (B) via a generalized linear mixed model (GLMM). The variation of the fixed effect estimates for each replicate (squares) and all replicates (diamonds) are shown as forest plots (95% confidence interval, glmmADMB). The square size is proportional to the sum of midguts analysed in each replicate. * P ≤ 0.05, ** P ≤ 0.01, ***P ≤ 0.001, **** P ≤ 0.0001 and ns not significant.

### *P. falciparum* transmission blocking assays

We assessed the effect of these modifications on mosquito infection with the *P. falciparum* NF54 strain by feeding transgenic female mosquitoes on *in vitro* cultured gametocytes using a standard membrane feeding assay (SMFA)(47). The number of oocysts in the midgut of wild type and the homozygous transgenic strains was quantified 7 days pbf (Figure 5). We did not observe a significant effect on oocyst intensity or prevalence in strains ScoG^CFP^-AP2 and Aper1-Sco^GFP^ (Figure 5A). This indicated that the observed reduction in host gene expression that occurs in both these strains had no measurable effect on transmission. Interestingly, strain Sco^GFP^-CP showed a significant reduction in infection prevalence (Figure 5A), whereas strain Sco-CP on the contrary, showed a substantially enhanced level of infection by both measures (Figure 5B). It had previously been shown, that expression of two *A. gambiae* carboxypeptidase B genes is upregulated upon *P. falciparum* infection and that antibodies against one of them blocked parasite development (48). Similarly, antibodies against the *A. stephensi* carboxypeptidases A and B significantly reduced the oocyst number of *P. berghei* and *P. falciparum*, respectively (49, 50). The strong decrease of CP mRNA expression in strain Sco^GFP^-CP could partially explain the observed effect. Strains ScoG-AP2 and Aper1-Sco both showed a highly significant reduction in *P. falciparum* oocysts (Figure 5B) in both infection prevalence and oocyst intensity lending support to the anti-malarial effect of Scorpine against NF54 parasites under these conditions.

### Analysis of non-autonomous gene drive induced by intronic gRNAs

Finally, we sought to determine whether the modified alleles of the 3 mosquito genes functioned as gene drives in the germline when provided with a source of Cas9 in *trans* (Figure 6). We assessed homing efficiency of each gRNA under the control of the U6 promoter within the intron, by crossing the Sco^GFP^-CP, ScoG^CFP^-AP2 and Aper1-Sco^GFP^ strains to a vasa-Cas9 strain (E. Marois, unpublished). Transhemizygote individuals were subsequently crossed to the wild type and the rate of fluorescent progeny was recorded (Figure 6). From this analysis, from which we excluded progeny that had also inherited Cas9 linked to the 3xP3-YFP marker (Figure 6 – figure supplement 1), we predicted a homing rate of 97.17% and 96.87% for Sco^GFP^-CP and Aper1-Sco^GFP^ respectively, with slightly higher rates in females compared to males (Table S5). Strain ScoG^CFP^-AP2 showed homing efficiency of 81.83%. Although this locus showed higher variability around the ATG insertion site in the Ag1000G dataset (51), no SNPs had been detected in 24 sequenced individuals from the G3 lab colony (52) suggesting that the lower homing rate was unlikely to be related to pre-existing resistant alleles.

**Figure 6.**
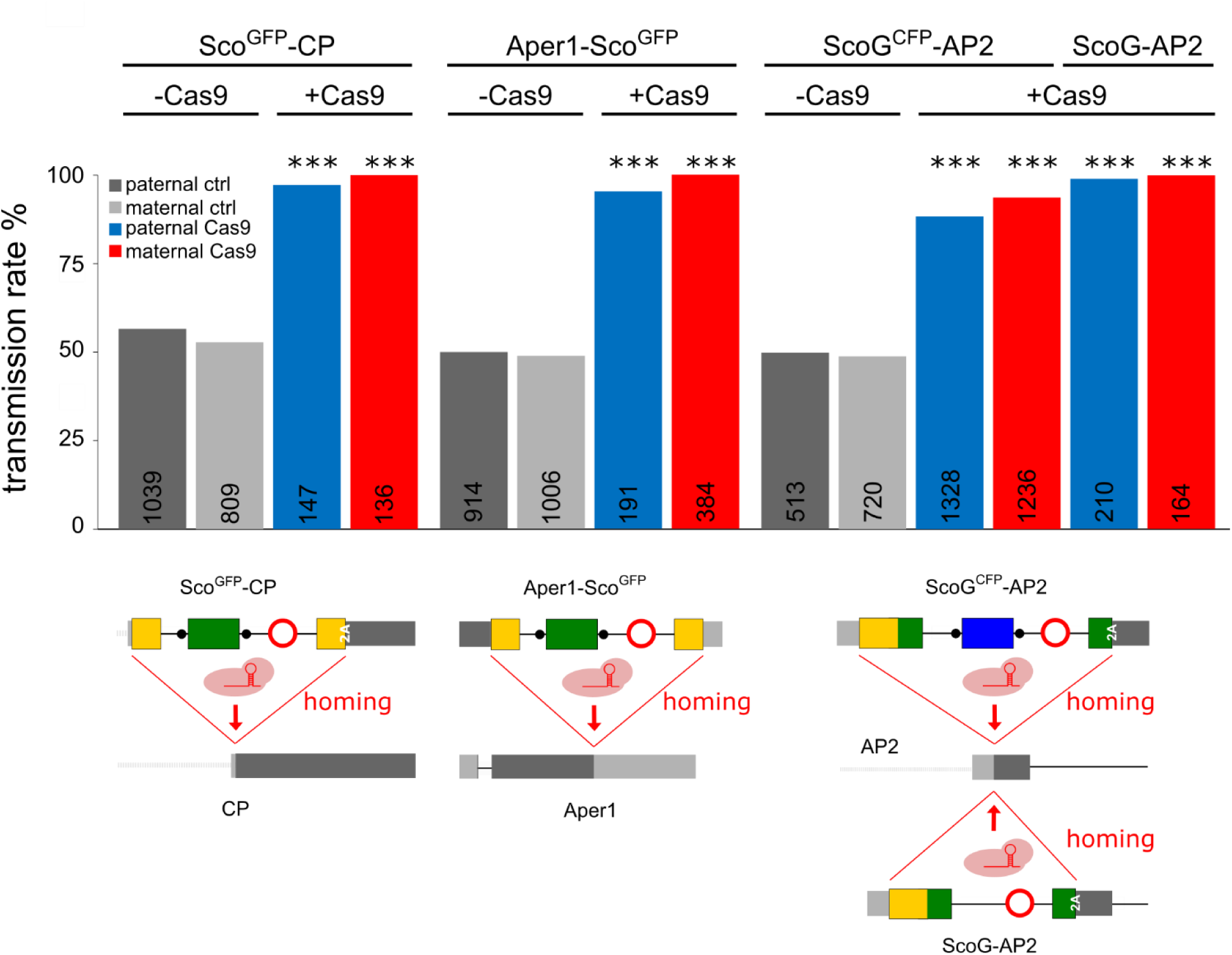
Assessment of non-autonomous gene drive of the modified host genes. Homozygous individuals of strains Sco^GFP^-CP, ScoG^CFP^-AP2 and Aper1-Sco^GFP^ or the markerless strain ScoG-AP2 were crossed to the vasa-Cas9 strain to assess the homing potential induced by the intronic guide RNAs. As a control, hemizygous individuals lacking Cas9 were crossed to WT. In each case homing was measured by the rate of fluorescent larvae recorded in the progeny with the exception of the markerless strain ScoG-AP2 where it was assessed via PCR genotyping of the progeny. The total number of individuals analysed is indicated for each column. All comparisons to the control crosses were significant (p < 0.0001, Chi-square test). The data is found in Supplementary Tables 4 and 5. The following figure supplements are available for figure 6. Figure supplement 1: Crossing schemes for assessing the homing rate. Figure supplement 2: Analysis of rearrangements in ScoG^CFP^-AP2 following Cas9 cleavage. Figure supplement 3: Genomic PCR sequencing of non-fluorescent individuals.

To better understand this finding, we analysed 41 non-fluorescent individuals obtained from the cross of Cas9,ScoG^CFP^-AP2 with wild type mosquitoes by PCR genotyping (Figure 6 – figure supplement 2A). For 29 out of 41 individuals analysed by primers spanning the entire locus, we detected a 2kb product in addition to the wild type amplicon. Sequencing these PCR products suggested rearrangements that are likely the result of homologous recombination between the GFP and CFP coding sequences (Figure 6 – figure supplement 2B). The remaining individuals showed wild type configuration at the guide RNA target site (Figure 6 – figure supplement 3A). We also sequenced the gRNA target region of all 9 non-fluorescent individuals obtained from the experiment with Aper1-Sco^GFP^ and observed no modified alleles or alleles corresponding to pre-existing SNPs within the gRNA (Figure 6 – figure supplement 3B).

We next sought to compare this result to the homing of a corresponding minimal genetic modification of the same locus. Homozygous strain ScoG-AP2 was crossed to the vasa-Cas9 strain and transhemizygotes were crossed to wild type individuals. For this experiment, where homing could not be tracked via a fluorescent marker, the resulting progeny was assessed via PCR genotyping with primers amplifying the Scorpine coding sequence. We recorded near complete homing with 372 out of 374 individuals inheriting the ScoG-AP2 allele. In summary, the strains described here can achieve near optimal homing rates of these effector cassettes in combination with a vasa driven Cas9 strain. We detected no events indicative of repair by non-homologous end joining.

## Discussion

We modified three different genomic loci of *Anopheles gambiae* via precise CRISPR/Cas9 homology-directed integration and subsequently removed the fluorescent transformation marker cassettes using Cre recombinase mediated excision. A range of methods for the generation of markerless, genetically modified organisms have been developed in plants (53). The presence of such marker genes in genetically modified plants, and subsequently in food, feed and the environment, are of concern and thus subject to special government regulations in many countries. With the view that gene drive strains that are currently being developed could eventually be deployed in the field and will have to undergo a stringent regulatory pathway and also that marker genes are known to induce negative fitness costs (54, 55), the generation of minimal genetic modifications is paramount for moving gene drives towards application. Combinations of multiple gene drives and multiple fluorescent markers would not only compound such costs but also reduce the usefulness of such markers. Furthermore, the predictive power of marker genes in the field will be low as gene drives can decouple from any linked marker cassettes within one generation (6, 56). We show for the first time how markerless gene drive traits can be constructed in the malaria mosquito and demonstrate multiple approaches to achieve this goal.

We expected that upon removal of the marker the remaining minimal modifications, i.e. the antimicrobial peptide Scorpine harbouring a small artificial intron would not significantly affect the function of the host genes. Unexpectedly, all strains including those carrying the entire fluorescent marker genes were found to be homozygous viable and fertile and showed no noticeable fitness cost during standard rearing conditions. RNA expression analysis indicated that host gene expression had indeed been disrupted or severely reduced in all 3 strains carrying the full insert including the fluorescent marker irrespectively of whether the host gene had been modified at the 5’ or 3’ end. The lack of a severe fitness phenotype could be explained by a certain redundancy in the set of digestive enzymes, proteases, phosphatases and peritrophins expressed in the mosquito midgut, at least under standard rearing conditions, including a high-quality source of human blood. Alternatively, splicing out of the entire intron including the marker cassette could occur at low levels. Our strains Sco^GFP^-CP, ScoG^CFP^-AP2 and Aper1-Sco^GFP^, thus will be useful tools in studying the function of the genes CP, AP2 and Aper1 under other conditions. Following the removal of the marker gene, we found by RNA expression analysis that exact splicing of the artificial intron restored expression of the host genes carrying now only a minimal transgene insert. The intron derives from the *Drosophila melanogaster* homeobox transcription factor *fushi tarazu* (*ftz*), essential for early embryonic development, and we have separately shown that a full gRNA transgene can be hosted within this intron (A. Nash et al. 2020, in preparation). Our data here show similarly that the *ftz* intron can accommodate the *A. gambiae* U6 promoter and the gRNAs for gene drive and remains functional in mosquitoes. The splicing was precise, in the context of Scorpine (AAG-intron-GG) or when inserted within Scorpine::GFP (AGG-intron-AG).

When we provided a source of Cas9, all gRNA transgenes were able to induce high rates of gene drive of the modified loci. In the case of the AP2 locus we found that the minimal, markerless locus ScoG-AP2 outperformed ScoG^CFP^-AP2. We observed aberrant homing events in the latter strain, which is likely the result of recombination between the GFP and CFP coding sequences. This highlights that simple genetic modifications are advantageous when it comes to homing but also that exogenous sequences of more complex constructs may not behave as expected. The interaction of multiple gene drives (sharing homologous sequences such as marker genes, promoters or terminators) could thus lead to unintended outcomes in the field. The regulatory regions of the target genes CP and Aper1 had previously been used for mosquito transgenesis (10, 11). In contrast, AP2 chosen due to differential expression in the female gut has not been described before. Together, our data on all 3 genes suggest, that co-opting the regulatory elements of endogenous loci directly without prior labour-intensive characterization is a viable approach. Since there is a dearth of promoters for many infection-relevant tissues (e.g. salivary glands, hemocytes, fatbody) we propose that this characterization step could be avoided, and effector molecules directly incorporated into mosquito gene loci and mobilized by non-autonomous drive. Existing RNA expression datasets or enhancer-trap screens could provide a starting point for the identification of suitable genes (57). However, our analysis revealed that modifications of endogenous genes could also have unforeseen consequences. In the case of the zinc carboxypeptidase A1 (CP) we found that modifications of the host gene are possibly directly altering infection outcomes. The reduced infection level observed for strain Sco^GFP^-CP which does not express CP (Figure 3C) is consistent with reports on other carboxypeptidase genes (48-50). However, strain Sco-CP, which features an N-terminal modification of CP and restored expression of CP displayed a direct positive effect on infection levels suggesting that the CP locus may not be a suitable host gene for gene drives. An alternative explanation could be that fixed genetic polymorphisms linked to the modified locus in this inbred strain could influence parasite infections.

We observed a substantial reduction in *P. falciparum* transmission in strains ScoG-AP2 and Aper1-Sco. Scorpine was chosen for our study as a prototypical effector molecule as it had previously shown to reduce *Plasmodium* survival (33-36) when expressed via different routes in the mosquito midgut. We do not know how the midgut environment containing digestive enzymes and proteases affects the stability of the lysine-rich Scorpine and detection of even the host genes in these samples proved difficult. However, modifications of the host genes AP2 and Aper1 did not appear to alter infection levels on their own, thus making these two loci suitable integration sites to test a range of other effector molecules. Such an approach would allow for direct comparisons between different classes of effectors either by secretion into the gut or by anchoring these molecules in the peritrophic matrix.

The most significant knowledge gap, when it comes to transmission blocking modifications of transgenic mosquitoes, is to what degree any effects observed with lab strains of *P. falciparum* would be reproducible using circulating parasites isolated from patient blood or would hold up under actual field conditions which could wildly differ from standard laboratory experiments. The strains we have described could be valuable tools to attempt to answer this question. The ScoG-AP2 and Aper1-Sco loci are both incapable of autonomous gene drive and have been established and maintained, unlike suppression drives where this is impossible, as true-breeding strains. They show a significantly reduced transmission potential under laboratory conditions using *P. falciparum* NF54. Importantly, the regulatory burden for importing or generating such non-autonomous effector strains would be expected to be much lower than strains capable of full gene drive and hence, simplify advanced stages of testing against polymorphic isolates of the *P. falciparum* parasite in the endemic setting without the need for strict containment or geographical isolation. When combined with other strains capable of autonomous Cas9 gene drive, whether designed for replacement or suppression, these strains could, without any further genetic modification, be deployed to contribute to reducing malaria transmission by mosquito populations in the field.

## Materials and Methods

### Plasmids & primers

For details on the generation of the donor plasmids pD-Sco-CP, pD-ScoG-AP2 and pD-Aper1-Sco (full plasmid sequences are provided in Supplementary file 1) see Supplementary Materials and Methods, for primers and intermediate plasmids see Tables S2 and S3 respectively.

### Microinjections and establishment of transgenic strains

Plasmids were isolated with the ZymoPURE II Plasmid Maxiprep kit (Zymo Research) and 300 ng/µL of the donor-plasmid and the p155-helper-plasmid (6) were microinjected into 30-45 minutes old eggs as described previously (58). Hatching larvae were screened for transient expression of the fluorescent marker and the adults crossed to wild type. The G1 offspring were screened for the presence of the GFP (Sco^GFP^-CP, Aper1-Sco^GFP^) or CFP (ScoG^CFP^-AP2) markers and we selected against red fluorescence in the eyes, in order to exclude individuals with possible plasmid backbone integration events, which was only observed in the case of ScoG^CFP^-AP2. G1 transgenic founder adults were backcrossed to WT and following egg laying, they were sacrificed and genomic DNA isolated. We performed PCR over the 5’ and 3’ insertion points using one primer binding within the construct and another primer binding the flanking genome sequence beyond the homology arms. Genomic DNA was obtained from single adults via incubation in 100µL of 5% w/v Chelex beads (BioRad Inc.) at 55°C, and all genotyping was performed with RedTaq Polymerase (VWR). Since for ScoG^CFP^-AP2 we obtained more than 100 G1 transgenic founder individuals, 12 single crosses were prepared in cups and offspring pooled after the parents were genotyped by sequencing.

### RT-PCR and qPCR

WT and transgenic mosquitos were dissected 3h after blood-feeding and tissue from 10 to 15 guts homogenized with glass beads for 30s at 6,800rpm in a Precellys 24 homogenizer (Bertin) in Trizol. RNA was extracted with the Direct-zol RNA Mini-prep kit (Zymo Research) and converted into cDNA with the iScript gDNA Clear cDNA Synthesis Kit (Bio-Rad). RT-PCR was performed with RedTaq Polymerase (VWR) with primers 51-Scorpine-F and 117-CP-ctrl-R on Sco-CP (1267bp), 160-Sco-probe-F and 6-F2A-R on ScoG-AP2 (1089bp) and 105-Per-locus-F and 162-Sco-probe-2-R on Aper1-Sco (533bp). The products were subsequently evaluated via gel-electrophoresis and sequencing (Genewiz). qPCRs were performed in triplicates with the Fast SYBR Green Master Mix (Thermo Scientific) on an Applied Biosystems 7500 Fast Real-Time PCR System. Expression was normalized to the S7 reference gene and analysed using the ΔΔCt method. The following primer pairs were used for the qPCRs: 272-qPer1-F1 and 273-qSco1-R2, 274-qPer1-F and 275 qPer1-R, as well as 214-qSco1-F2 and 215-qSco1-R.

### Mass-spectrometry

Following dissection, 25 non-blood-fed guts of strains Sco-CP or Aper1-Sco were put in 50µL CERI reagent from the NE-PER™ Nuclear and Cytoplasmic Extraction kit (Thermo Fisher Scientific), homogenised with a motorized pestle and cytoplasmic proteins were extracted according to the manufacturers protocol. The supernatant was filtered through a 100kDa or 50kDa Amicon Ultra-0.5 Centrifugal Filter Unit for Sco-CP and Aper1-Sco, respectively. Trypsin-digest and LC-MS/MS was performed at the Advanced Mass Spectrometry Facility at the University of Birmingham.

### Fitness assays

For fecundity and fertility assays, single blood-fed females were transferred to cups containing water, lined with filter paper. Females that failed to lay eggs or produce larvae, were dissected and excluded from the analysis if no sperm was detected in their spermatheca. Eggs and L1 larvae from at least 10 technical replicates were counted and the data from 3 biological replicates were pooled. All biological replicates passed the Shapiro-Wilk normality test. To determine the pupal sex ratio, between 100 and 140 L1 larvae per tray were reared to the pupal stage and sexed. Data from 3 biological replicates was pooled and analysed for deviations from the expected 1:1 sex ratio via the Chi-square-test.

### *Plasmodium falciparum* standard membrane feeding assay

Infections of mosquitoes using the streamlined Standard Membrane Feeding Assay were performed as described previously (47). Briefly, mosquitoes were fed for 15 min at room temperature using an artificial membrane feeder with a volume of 300–500µl of mature *Plasmodium falciparum* (NF54) gametocyte cultures (2-6% gametocytaemia). Afterwards, mosquitoes were maintained at 26°C with 70–80% relative humidity. For 48 hours after the infective meal, mosquitoes were deprived of light and starved without fructose, in order to eliminate the unfed mosquitoes. Midguts were dissected after 7 days. Statistical significance was calculated using the non-parametric Mann-Whitney test for oocyst load (infection intensity) and the Chi-square test for oocyst presence (infection prevalence) with GraphPad Prism v7.0. The generalized linear mixed model (GLMM) was used to determine statistical significance of oocyst infection intensity for each independent biological replicate. GLMM analyses were performed in R (version 1.2.5019) using the Wald Z-test on a zero-inflated negative binomial regression (glmmADMB). The different strains were considered as covariates and the replicates as a random component. Fixed effect estimates are the regression coefficients.

### Assessment of the homing rate

Homozygous individuals of strains Sco^GFP^-CP, ScoG^CFP^-AP2 and Aper1-Sco^GFP^ were crossed to the vasa-Cas9 strain carrying the 3xP3-YFP marker (chromosome 2) and the offspring were screened for the presence of red fluorescence (Figure 4 – figure supplement 1). Subsequently, the transhemizygotes were sexed and crossed to G3 wild types. Offspring deriving from the ScoG^CFP^-AP2 cross were screened for the presence of CFP, whereas progeny deriving from Sco^GFP^-CP and Aper1-Sco^GFP^ were selected based on the absence of the RFP and the presence of the GFP marker. Progeny from the cross of ScoG-AP2/Cas9 with wild types were screened by PCR. As controls, hemizygous individuals of each strain were crossed to WT, and the rate of fluorescent larvae was calculated. For the calculation of the homing rate e, we used the individuals negative for the effector (E_neg_), the number n and the Mendelian distribution of 50% as a baseline, as follows: e = (n*0.5 – E_neg_) / (n*0.5) *100.

## Acknowledgments

We thank Alexander Nash, George Avraam, Olivia Bates, Claudia Wyer, Louise Marston, Roberto Galizi, Andrew Hammond, Carla Siniscalchi and Eric Marois, as well as Jinglei Yu from the Advanced Mass Spectrometry Facility at the University of Birmingham. The work was funded by the Bill and Melinda Gates Foundation grant OPP1158151 to N.W. and G.K.C

