## Supplementary Information for "Converting endogenous genes of the malaria mosquito into simple non-autonomous gene drives for population replacement"

### Supplementary Materials and Methods

**Plasmid construction**

The two intermediate plasmids pI-Scorpine-GFP and pI-Scorpine were generated first to allow insertion of different gRNAs. We synthesized the Scorpine coding sequences, optimized for codon usage in *Anopheles gambiae* and retained the endogenous secretion signal from Scorpine. The GFP containing the intron, based on the *Drosophila melanogaster fushi tarazu* (ftz) intron, was amplified from pQUAST 3xP3 CFP (A. Nash et al. 2020, in preparation) and the U6 promoter was exchanged for the *A. gambiae* U6 promoter from p165 (1). The EGFP marker module within pI-Scorpine was amplified from p163 (1). The intron within Scorpine was inserted between the nucleotides AAG and G to promote splicing (2). We used the Drosophila splice predictor (3) to remove cryptic splice sites. A furin cleavage site and the F2A peptide (4) was generated via annealing long oligos (53-F2A-long-F and 54-F2A-long-R). Some of the initial fragments were pre-fused via overlap extension PCR in order to minimize the amount of parts to be fused for the final Gibson assembly.
The guide-RNAs for the 3 target loci were generated via the annealing of oligonucleotides 39-CP-gRNA59-F and 40-CP-gRNA59-R, 57-AP2-gRNA53-F and 58-AP2-gRNA53-R, as well as 75-Per1-gRNA48-F and 76-Per1-gRNA48-R. They were then inserted into the intermediate plasmids through the BbsI restriction site via Golden gate cloning. Subsequently, each effector-cassette hosting the corresponding gRNA was PCR-amplified from the resulting intermediate plasmids. For the donor plasmids pD-ScoG-AP2 and pD-Sco-CP, the cassette was amplified with primers 51-Scorpine-F and 6-F2A-R, and for pD-Aper1-Sco with primers 77-Arg-GSG-Arg-Scorp-F and 97-Scorpine-STOP-R.
The 5’ and 3’ homology arms of approximately 800 bp were amplified from *A. gambiae* G3 genomic DNA. All amino acid changes detected in protein sequences were confirmed via the variation data available in www.vectorbase.org, except the A23V amino acid change in the secretion signal of the CP CDS. Nevertheless, this variant was observed in the 24 individual G3 sequencing reads. Since the guide RNA for Aper1 does not overlap with the STOP codon, the 5 wobble bases of the 17 re-inserted bases were re-coded in order to destroy the guide RNA target in the homology arm.
The donor-plasmids were generated by assembling the homology arms, the cassette and the backbone containing an additional 3xP3-DsRed marker module. All plasmids were propagated in Agilent Sure cells in order to avoid recombination. For primers and plasmids see Tables S2 and S3.

**Establishment of pure breeding transgenic strains**

$Sco^{GFP}$-CP and $ScoG^{CFP}$-AP2 were rendered homozygous by setting up sibling crosses over two generations and letting the adults hatch single in cups and genotyping their left-over pupal cases by PCR over the entire locus to distinguish WT, heterozygotes and homozygotes. $Sco^{GFP}$-CP was genotyped with primers 99-CP-locus-F and 100-CP-locus-R, and $ScoG^{CFP}$-AP2 with 101-AP-locus-F and 102-AP-locus-R. In the case of Aper1-$Sco^{GFP}$, the transgene was crossed to a vasa-Cas9 strain (E. Marois, unpublished), and kept in this background for several generations to allow for homing to occur. The YFP marker is also visible in the green channel and was removed via screening for red fluorescence at a wavelength of 563 nm (Filter 630/30 nm emission, 595 nm dichroic) on a Nikon inverted microscope (Eclipse TE200). Afterwards, crosses of 4 females to 4 males were prepared in cups and the parents were genotyped with 230-Per-short-F and 231-Per-short-R after egg laying. gDNA was isolated from pupae cases in 20µL dilution buffer of the Phire Tissue Direct PCR Master Mix kit (Thermo Scientific).

**Removal of the marker module using Cre recombinase**

Markerless strains were generated by removal of the fluorescent marker cassette through crossing to strain CS2 (5) expressing Cre recombinase driven by the vasa promoter and carrying a 3xP3-DsRed marker (herein referred to simply as Cre). The Cre transgene is on the 3rd chromosome and resides in a KIL background and contains an additional 3xP3-CFP marker (6). In the case of CP, the crossing scheme was initiated with homozygous $Sco^{GFP}$-CP individuals and offspring showing green fluorescence and red fluorescence for Cre were kept for setting up a siblings-cross (Figure 2B). The progeny from $Sco^{GFP}$-CP/Cre were screened against the presence of both markers and 88 larvae still containing the Cre transgene were discarded. There were no more larvae with GFP present, and hence, the efficiency of the Cre-mediated excision was 100%. 32 individuals were marker-less and were genotyped by PCR on pupal cases for homozygosity. 7 individuals were homozygous, 3 hemizygous, one WT and 18 PCRs had unclear results. A pure breeding strain was established immediately afterwards from 3 male and 2 female markerless founders.

In the case of Aper1, the crossing scheme was started with hemizygous Aper1-$Sco^{GFP}$ individuals and the offspring were screened for GFP and DsRed (Figure 2 – figure supplement 1A). After crossing them to the Cas9 strain, the progeny was screened against CFP to remove the Cre transgene but were kept in this Cas9 background over several generations, in order to make them homozygous. After removal of the Cas9 transgene, 13 final founders were confirmed via genotyping and sequencing over the lox-out, as well as over the 5´and 3´ insertion sites and no imprecise homing events were detected.

In the case of the AP2, we used homozygous $ScoG^{CFP}$-AP2 individuals to initiate the cross to the Cre strain which also features the 3xP3-CFP marker (Figure 2 – figure supplement 1B). Offspring exhibiting both blue and red fluorescence were crossed to the vasa-Cas9 strain and the progeny were screened for the presence of YFP and the absence of the CFP and DsRed markers, which would maintain the markerless transgene in a Cas9 background. One further generation was propagated without screening in order to let the cassette home. Subsequently, individuals were genotyped using pupal cases and homozygotes were crossed to homozygous $ScoG^{CFP}$-AP2 individuals. The offspring were screened against the presence of the YFP to select against Cas9. Finally, ScoG-AP2/$ScoG^{CFP}$AP2 mosquitoes were crossed and the progeny were screened against CFP. The majority of them was also genotyped after egg laying.

**Homing rate assessment**

20 $Sco^{GFP}$-CP;Cas9 females were crossed to 20 G3 WT males, and 20 $Sco^{GFP}$-CP;Cas9 males were crossed to 20 G3 WT females. 78 male Aper1-$Sco^{GFP}$/Cas9 were crossed to 120 WT females, and 68 female Aper1-$Sco^{GFP}$/Cas9 to 120 WT males. 44 male $ScoG^{CFP}$-AP2/Cas9 were crossed to 64 female G3 WT, 27 female $ScoG^{CFP}$-AP2/Cas9 were crossed to 57 male G3 WT, and an additional 14 females to 20 males. In order to evaluate the homing rate of the markerless AP2-line, 36 females of ScoG-AP2/Cas9 were crossed to 33 WT males and 71 offspring were screened by PCR for the presence of the construct. Another 13 females of ScoG-AP2/Cas9 were crossed with 60 WT males and 71 individuals of the progeny were screened by PCR. 24 males of ScoG-AP2/Cas9 were crossed to 55 WT females and 213 offspring were screened by PCR. gDNA was isolated with Chelex beads (BioRad) or the dilution buffer of the Phire Tissue Direct PCR Kit (Thermo Scientific) and PCRs were performed with primers 101-AP-locus-F and 102-AP-locus-R, and in case of doubt another PCR with primers 160-Sco-probe-F and 161-Sco-probe-R was performed. An additional 62

individuals from the Aper1-Sco$^{GFP}$/Cas9 males were screened by PCR with primers 230-Per-short-F and 231-Per-short-R.

### Supplementary Figures

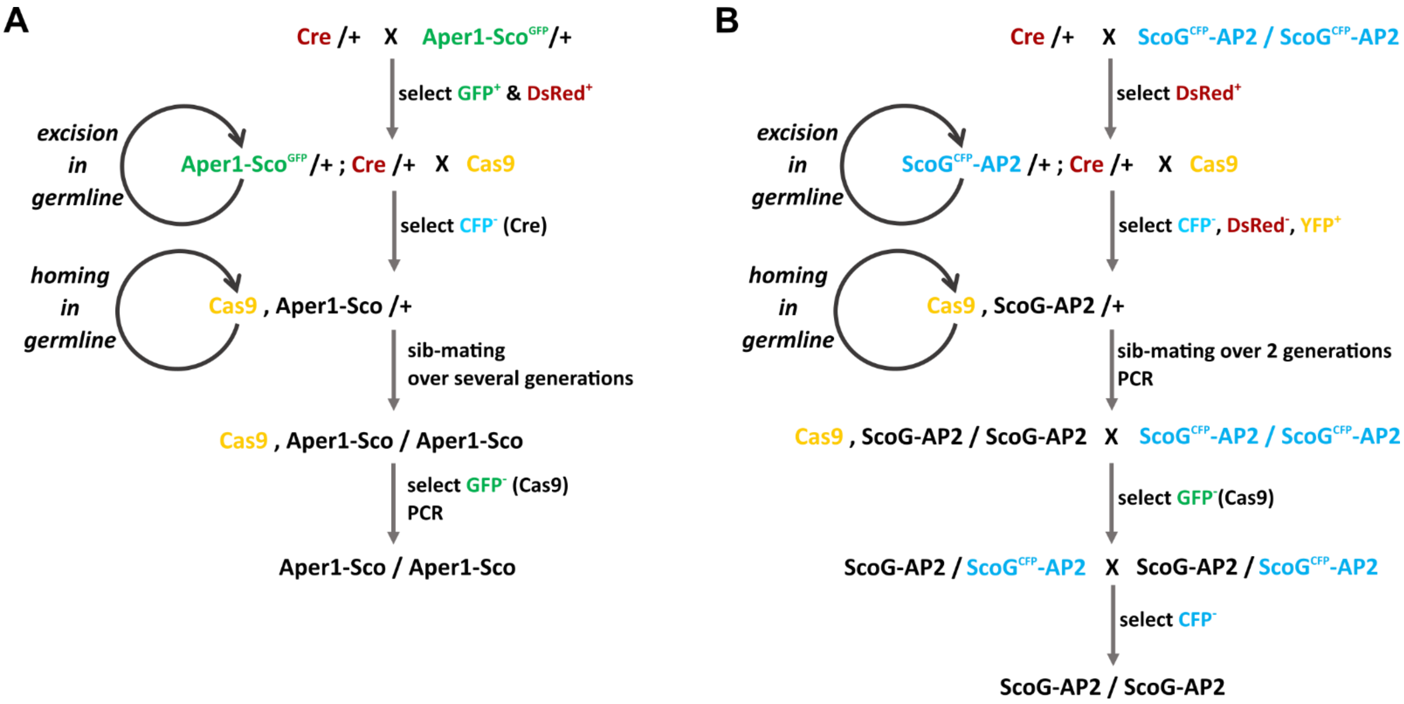


**Figure 2 – figure supplement 1: Crossing schemes used for the establishment of markerless strains.** The strains Aper1-Sco (A) and ScoG-AP2 (B) were rendered homozygous by first crossing to a Cre recombinase and then a Cas9 expressing strain (Eric Marois, unpublished) and by employing positive and negative selection via the fluorescent markers at each stage.

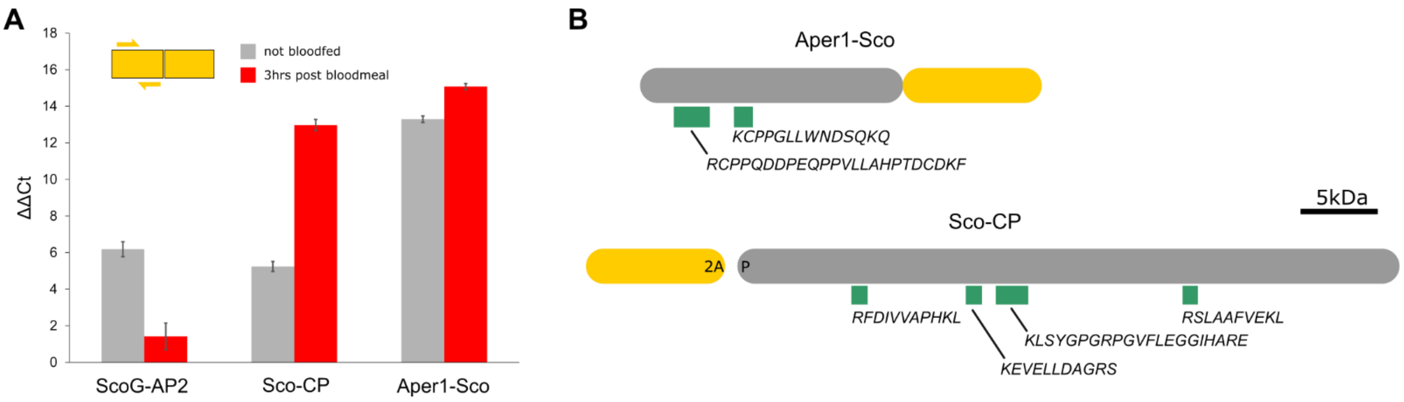


**Figure 3 – figure supplement 1: Expression levels and mass-spectrometry of co-opted host genes.**
**(A)** qPCR for the integrated Scorpine sequence was performed on cDNA from both non-blood-fed guts and guts dissected 3 hours after blood-feed of all 3 markerless transgenic strains. The primer-pair binding to Scorpine is indicated as yellow half arrows in the inset. Error bars indicate standard deviation from 3 technical replicates normalized to the S7 reference gene. G3 wild-type controls (blood-fed and non-blood-fed) run in parallel were negative (Ct mean values above 35).
**(B)** Peptides identified in mass-spectrometry that map to the endogenous host genes. Proteins were extracted from non-blood-fed guts of the homozygous transgenic lines Sco-CP and Aper1-Sco. Note that Sco-CP (but not Aper1-Sco) is expected to generate two protein products due to the presence of the 2A peptide. Green rectangles indicate high confidence peptides with amino acid sequences shown below.

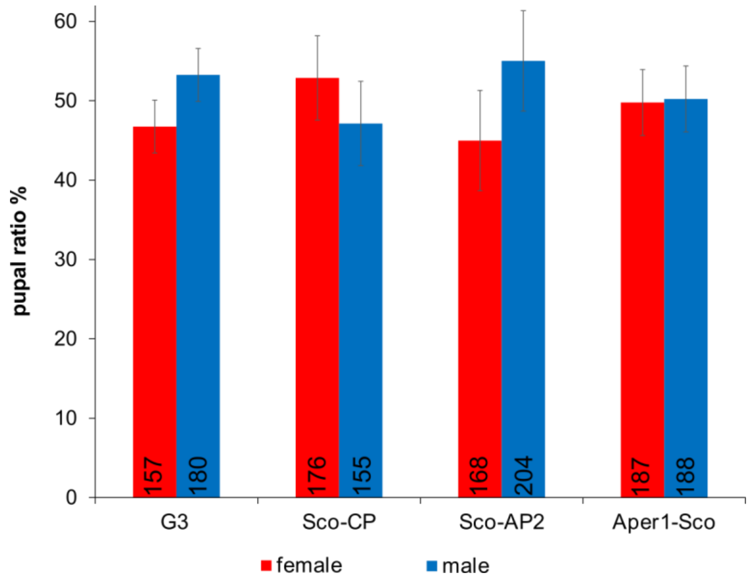


**Figure 4 - figure supplement 1: Pupal sex ratio.**
Pupal sex ratio of the homozygous markerless transgenic strains Sco-CP, Sco-AP2 and Aper1-Sco compared to the G3 wild-type control. No statistically significant deviation from an expected 1:1 sex ratio was detected (Chi-square-test). Error bars represent standard deviation from 3 biological replicates and the number of counted individuals is indicated (n).

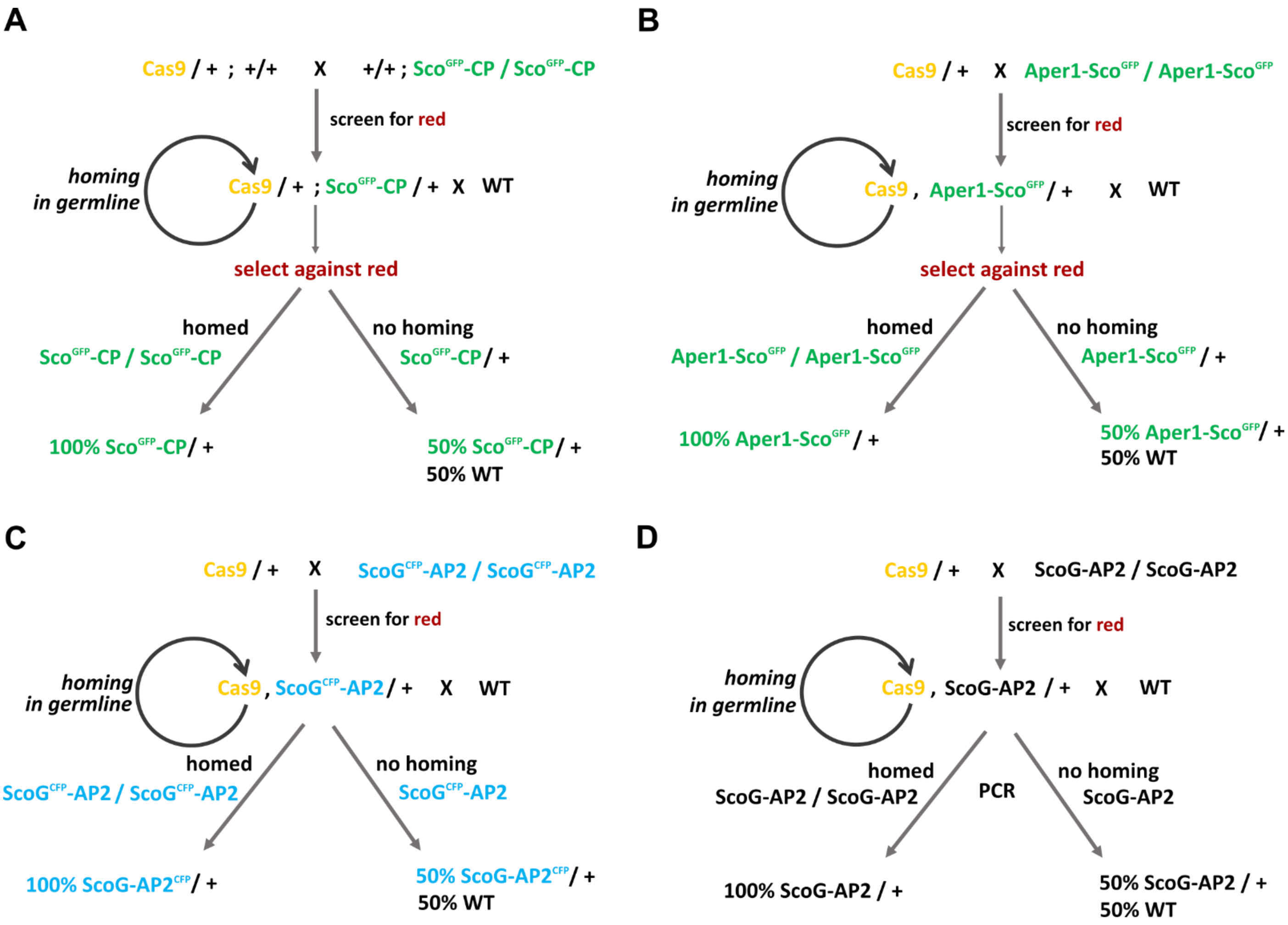


**Figure 6 – figure supplement 1: Crossing schemes for assessing the homing rate.**
The non-homing vasa-Cas9-line carrying a 3xP3-YFP marker (Eric Marois, unpublished) was crossed to the transgenics. If homing in the germline occurs, all progeny from the cross to WT will be heterozygotes. In the case of Sco$^{GFP}$-CP and Aper1-Sco$^{GFP}$, only the larvae without the Cas9 transgene can be considered for the assessment, due to an overlap of fluorescent markers in the GFP channel.

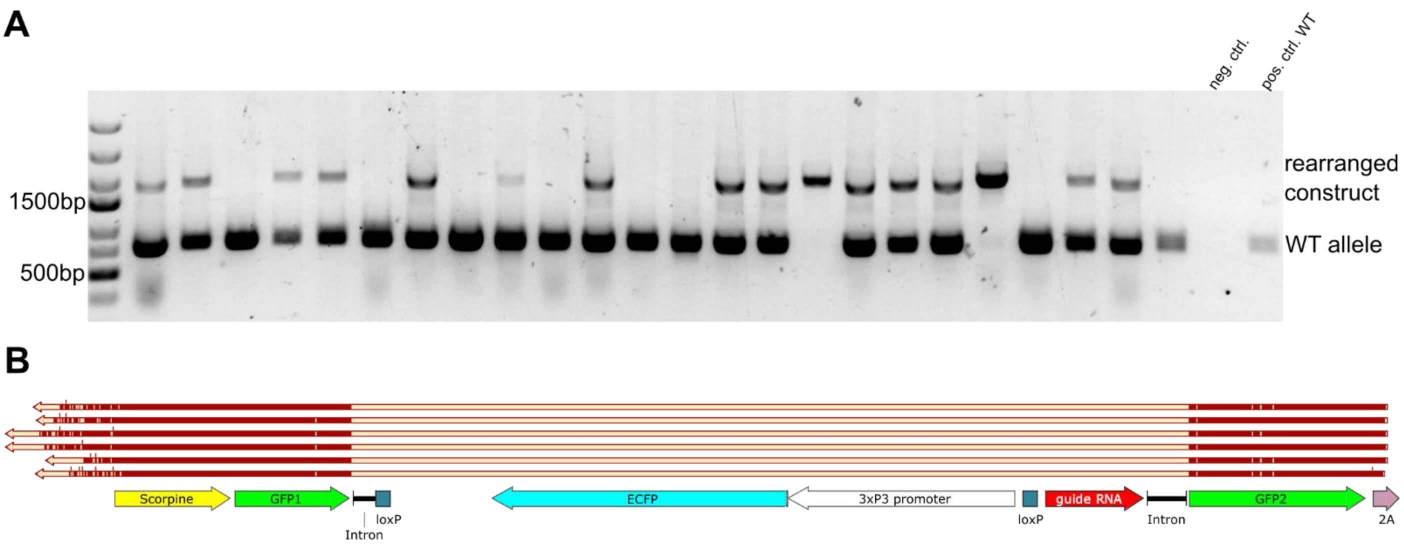


**Figure 6 – figure supplement 2: Analysis of rearrangements in ScoG$^{CFP}$-AP2 following Cas9 cleavage.** (A) PCR analysis of non-fluorescent offspring from the cross of ScoG$^{CFP}$-AP2/Cas9 to WT with primers spanning the locus showed that the majority of individuals carries an insertion but of reduced size.
(B) Sequencing result of the upper band identified in the PCR suggests the loss of the intron cassette as well as a recombination and sequence exchange between the GFP and CFP coding sequences.

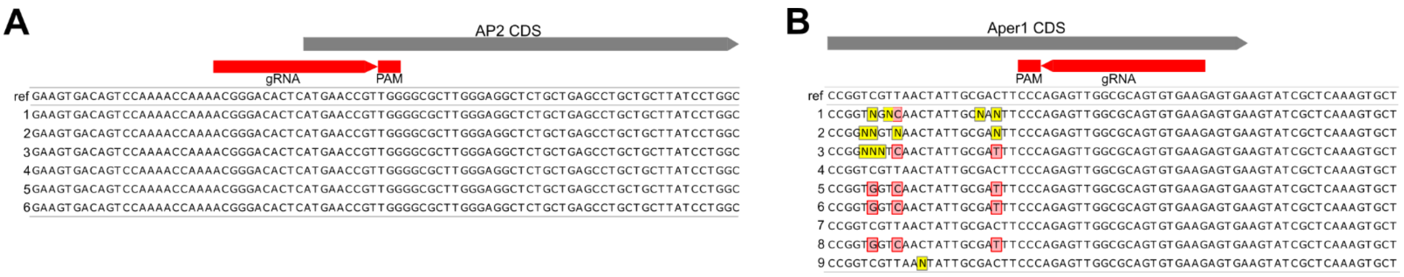


**Figure 6 – figure supplement 3: Genomic PCR sequencing of non-fluorescent individuals.**
The products of PCRs over the gRNA target sites, performed on 6 and 9 non-fluorescent individuals identified in the homing assay with ScoG$^{CFP}$-AP2 (A) and Aper1-Sco$^{GFP}$ (B) respectively, were sequenced. SNPs are indicated in red, and all of them coincide with wobble bases in the Aper1 coding sequence. N (in yellow) represent undetermined sequencing results.

### Supplementary Tables

#### Table S1: Guide RNA and target site characteristics.

| gene | AGAP | chrom. | gRNA-PAM | activity | off-target score | off-targets | MM | G3 | Ag1000G |
|---|---|---|---|---|---|---|---|---|---|
| CP | 009593 | 3R | AGCAAGCGGTCGATTGAAC**A-TG**G | 59 | 50 | 3 NC | 0/2/3 | 0 | no pass |
| AP2 | 006400 | 2L | ACGGGACACTC**ATG**AACCGT-TGG | 53 | 99 | 1 NC | 3 | 0 | 9 |
| Aper1 | 006795 | 2L | CTTCACACTGCGCCAACTCT-GGG | 48 | 98 | 4 NC | 3 | 0 | 4 |

The start codon of the target gene is indicated within the gRNA sequence (bold) and the protospacer adjacent motif (PAM) separated via a hyphen. All predicted off-target cleavage sites were found to be located in non-coding (NC) regions and the number of mismatches (MM) is indicated. The number of SNPs within the 24 individuals of the G3 strain and within the Ag1000G is indicated. The SNPs observed for CP in the Ag1000G did not pass the quality control.

### Table S2: Primers used in this study

| number | name | sequence |
|---|---|---|
| 5 | GFP-F | GTGAGCAAGGGCGAGGAGCTG |
| 6 | F2A-R | AGGGCCCGGGTTCGACTCCAC |
| 7 | SpeI-ori-F | ACTAGTCGCGTTGCTGGCGTTTTTCC |
| 8 | Amp-R | AACCGTGCGTTATTTTTCTAAATACATTCAAATATG |
| 9 | 3xP3-F | GAAAAATAACGCACGGTTCCCACAATGGTTAATTC |
| 10 | BB-SV40-R | CTCGAGAAGATACATTGATGAGTTTGG |
| 15 | U6Term-AflII-R | acgacggccagtcttaaggggacccacgcgtaaaaaaaagc |
| 26 | lox-AgU6-F | aagtgacacgacgccgattttgtatgcgtgcgcttgaag |
| 27 | AgU6-Bbs-R | AGGTCTTCTCGAAGACCCCagcagagagcaactccatttc |
| 28 | BB-scaffold-F | GGGTCTTCGAGAAGACCTgttttagagctagaaatagcaag |
| 29 | BB-lox-R | ttcaagcgcacgcatacaaaatcggcgtcgtgtcacttc |
| 30 | ori-F | tgcttttttttttccataggctccgcccccc |
| 31 | Amp-R2 | tcgcgttctacataccggtcgcggaacccctatttgttta |
| 32 | vas-AgU6-F | cacattttcgggcgcgcctttgtatgcgtgcgcttgaag |
| 33 | scaffold-ori-R | tatggaaaaaaaaaaagcaccgactcggtgccac |
| 39 | CP-gRNA59-F | tgctGAGCAAGCGGTCGATTGAACA |
| 40 | CP-gRNA59-R | aaacTGTTCAATCGACCGCTTGCTC |
| 41 | CP-HA5-F | ATCAATGTATCTTCTCGAGTGTTGATCGGGTTTGACACAAAG |
| 42 | CP-HA5-R | agaTCTTGGAGAAGTTCATgtTCAATCGACCGCTTGCTTGCAC |
| 43 | CP-HA3-F | TGGAGTCGAACCCGGGCCCTATGGTGCGATTAAACAGTGCAG |
| 44 | CP-HA3-R | ACGCCAGCAACGCGACTAGTATTGCTCGTGCCCTGCTCGGCCCA |
| 47 | between-F | attatgatctagagtcgcggccgc |
| 48 | between-R | ccgcgactctagatcataatcag |
| 50 | BB-Kozac-R | cgtcagcttcgagttcatggtggcgcagatctgttaacg |
| 51 | scorpine-F | ccaccatgaactcgaagctgacggccctg |
| 52 | scorpine-R | agctcctcgcccttgctcacTCCGG |
| 53 | F2A-long-F | aagCGCGCCAAGCGCGCCCCGGTGAAGCAGACGCTGAACTTCGACCTGCTGAAGCTGGC |
| 54 | F2A-long-R | CGGGTTCGACTCCACGTCGCCGGCCAGCTTCAGCAGGTCGAAGTTCAGCGTCTGCTTCA |
| 55 | GFP-furin-R | CCGGGGCGCGCTTGGCGCGcttgtacagctcgtccatgccgag |
| 56 | F2A-BB-F | CGGCGACGTGGAGTCGAACCCGGGCCCTtctagaggatctttgtgaaggaac |
| 57 | AP2-gRNA53-F | tgctGACGGGACACTCATGAACCGT |
| 58 | AP2-gRNA53-R | aaacACGGTTCATGAGTGTCCCGTC |
| 59 | AP2-HA5-F | TCAATGTATCTTCTCGAGATGCGAAACAAAGCAATCCACGAC |
| 60 | AP2-HA5-R | tcagcttcgagttcatggtggGAGTGTCCCGTTTTGGTTTTGGACTG |
| 61 | AP2-HA3-F | AGTCGAACCCGGGCCCTATGAACCGTTGGGGCGCTTGGGAGG |
| 62 | AP2-HA3-R | ACGCCAGCAACGCGACTAGTTCTACTGTTTCTGAACTTCTGCAATC |
| 69 | RGSG-Scorp-F | AgGAacgcGGATCCGGAggctggatcaacgaggagaagatcc |
| 70 | Per1-GFP-R | gcgATACTTCActtgtacagctcgtccatgccgag |
| 71 | Per1-HA5-F | TCAATGTATCTTCTCGAGACACTTTCTCACGAGCGGTCAGGGTG |
| 72 | Per1-HA5-R | ccTCCGGATCCgcgtTCcTCgCAtTGaGCtAAtTCaGGaAAGTCGCAATAG |
| 73 | Per1-HA3-F | tacaagTGAAGTATCGCTCAAAGTGCTCGAGCTG |
| 74 | Per1-HA3-R | GCCAGCAACGCGACTAGTAACACAGTACGGCGGAGTCTGTAAAGG |
| 75 | Per1-gRNA48-F | tgctGTTCACACTGCGCCAACTCT |
| 76 | Per1-gRNA48-R | aaacAGAGTTGGCGCAGTGTGAAC |
| 77 | Arg-GSG-Arg-Scorp-F | gcGGATCCGGAcgcggctggatcaacgaggagaag |
| 78 | GFP-R | cttgtacagctcgtccatgccgag |
| 85 | Scorp-F2A-F | gctgtcgtacCGCGCCAAGCGCGCCCCGGTGAAGC |
| 86 | intron1-Scorp-R | tgatacttaccttctcgcccgacgtctggcagtg |
| 87 | Scorp1-intron-F | gcgagaaggtaagtatcacacacgattaac |
| 88 | SV40-loxP-R | atgtatcttaagctataacttcgtataatgtatg |
| 89 | loxP-SV40-F | ttatagcttaagatacattgatgagtttgg |
| 90 | 3xP3-intron-R | tctagaactagtggatcttaattaaccacaatggttaattc |
| 91 | marker-lox-F | aagatccactagttctagagcggataac |

| | | |
|---|---|---|
| 92 | intron2-Scor2-R | gccgcacttgcacttcgtgccgtggcagtagccctgtaagcataagcaaagaaaaaatg |
| 93 | CP-HA5-Scorp-R | tcagcttcgagttcatggtggTCAATCGACCGCTTGCTTGCACGAAC |
| 95 | Per1-HA5-Arg-GSG-R | agccgcgTCCGGATCCgcgtTCcTCgCAtTGaGCtAAtTCaGGaAAGTCGCAATAG |
| 96 | Per1-HA3-scorp-F | acgccgctgtcgtacTAAAGTATCGCTCAAAGTGCTCGAGCTG |
| 97 | Scorpine-STOP-R | TTAgtacgacagcggcgtgccgcacttgcacttcgtgccgtggcagtagcc |
| 98 | Scorp-F2A-long-F | acgaagtgcaagtgcggcacgccgctgtcgtacCGCGCCAAGCGCGCCCCGGTGAAGC |
| 99 | CP-locus-F | ATTATCGGAAATTCTCCACAAAGCG |
| 100 | CP-locus-R | TCATGGTGCGATTCTGATGCGATCG |
| 101 | AP-locus-F | ATCAGACTTTAAGCCTTTGTCGTC |
| 102 | AP-locus-R | ACCGTTCGCGATTCAGCAGTACAGG |
| 105 | Per1-locus-F | TGTCCACCCGGGTTGCTGTGGAATG |
| 106 | Per1-locus-R | AACTGTAAATGATATAACACTATCC |
| 109 | CP-takara-F | CCTCTAATACGACTCACTATAggAGCAAGCGGTCGATTGAACAGTTTAAGAGCTATGC |
| 110 | AP-takara-F | CCTCTAATACGACTCACTATAggACGGGACACTCATGAACCGTGTTTAAGAGCTATGC |
| 112 | Per1-takara-F | CCTCTAATACGACTCACTATAggTTCACACTGCGCCAACTCTGTTTAAGAGCTATGC |
| 116 | CP-ctrl-F | AAATACATGACTGAGGACCAAATTCC |
| 117 | CP-ctrl-R | TCCACAAACGCAGCCAGCGAACGGGTC |
| 118 | AP-ctrl-F | TCTCACGCTATCCCGCCCAATTAAGC |
| 119 | AP-ctrl-R | TTCTTTCAGTTTCCACCTGTCCAACC |
| 122 | Per1-ctrl-F | TTTCATTAGTGTGGACGCGGGAAC |
| 123 | Per1-ctrl-R | TCGACCGGAAATGGTAGAGTTCGGTCG |
| 128 | STOP-linker | TAAgTAAcTAAcataacttcgtatagcatac |
| 129 | scaff-U6term-R | acaaaaaagcaccgactcggtgccac |
| 160 | Sco-probe-F | ATGAACTCGAAGCTGACGGCCCTG |
| 161 | Sco-probe-R | ACGTCTGGCAGTGCTTCTCGCAGTTGC |
| 162 | Sco-probe-2-R | tacgacagcggcgtgccgcacttg |
| 212 | q-Sco1-F | agctgacggccctgatcttcc |
| 213 | q-Sco1-F1 | atgggcaacacggtgctgg |
| 214 | q-Sco1-F2 | actgcggctggatcaacgagg |
| 215 | q-Sco1-R | tcgcagttgcccagcatgtcc |
| 216 | q-Sco-int-R | tggcagtagcccttctcgcc |
| 217 | q-Sco2-R | TTGGCGCGgtacgacagcg |
| 218 | q-CP-F | AACGAACAAGCGCGCCAAGG |
| 219 | q-CP-R | ATTCTGATGCGATCGTCCTGCG |
| 220 | q-AP-F | TGAACCGTTGGGGCGCTTGG |
| 221 | q-AP-R | AGCTCGTGGGCCATATGCTTCC |
| 222 | q-GFP1-F1 | atctgcaccaccggcaagc |
| 223 | q-GFP-int-R | aagatggtgcgctcctggacg |
| 224 | q-GFP1-F2 | taccccgaccacatgaagcagc |
| 225 | q-GFP2-R | tcgccctcgaacttcacctcgg |
| 226 | q-linker-F | tcgtacGGATCCGGAgtgagc |
| 227 | q-F2A-R | AGTTCAGCGTCTGCTTCACCG |
| 228 | q-S7-F | TTCTCCGGCAAGCACGTCG |
| 229 | q-S7-R | ACGCTTGCCGACCACCTCC |
| 230 | Per-short-F | ATCCCGACCATATGGTGTACATTCC |
| 231 | Per-short-R | ACACGAGAAGAAACCCCCATTCC |
| 245 | q-Sco-int-F | tcgggcgagaagggctactgc |
| 246 | q-CP-R2 | AGCCGGCTGCACTGTTTAATCG |
| 270 | q-CP-F1 | ACGAGATTGCCGAGGCGAC |
| 271 | q-CP-R3 | AGCTCCAGTCAACGCTCGACC |
| 272 | q-Per1-F1 | TTGAGCTGGAGCAGACGTGCC |
| 273 | q-Sco1-R2 | tgttgcccatgcgctcgtcg |
| 274 | q-Per1-F | AAACCGTCACCCAACTGTCCG |
| 275 | q-Per1-R | TCCAGCTCAACGCCGTACGG |

**Table S3: Plasmids used in this study**

| plasmid | type | R | marker | notes |
|---|---|---|---|---|
| pQUAST 3xP3 CFP | template | Amp | 3xP3-CFP | A. Nash 2020, in preparation |
| p163-pattP-P3GFP | template | Amp | 3xP3-EGFP | Hammond et al. 2016 |
| p165-pvasa-hCas9-U6 | template | Chlor | 3xP3-RFP | Hammond et al. 2016 |
| p155-pattB-CFP-vas2-hCas9 | helper | Amp | 3xP3-CFP | Hammond et al. 2016 |
| pI-Scorpine-GFP | intermediate | Amp | 3xP3-CFP | intron in GFP, longer 3xP3 promoter |
| pI-Scorpine-GFP-AP2 | intermediate | Amp | 3xP3-CFP | |
| pD-ScoG-AP2 | donor | Amp | 3xP3-CFP | at ATG, F2A |
| pI-Scorpine | intermediate | Amp | 3xP3-EGFP | intron in Scorpine, shorter 3xP3 promoter |
| pI-Sco-CP | intermediate | Amp | 3xP3-EGFP | |
| pI-Sco-Per1 | intermediate | Amp | 3xP3-EGFP | |
| pD-Sco-CP | donor | Amp | 3xP3-EGFP | at ATG, F2A |
| pD-Aper1-Sco | donor | Amp | 3xP3-EGFP | at STOP, no F2A |

R: antibiotic resistance to Ampicillin (Amp) or Chloramphenicol (Chlor).

### Table S4: Transmission rate of control-crosses without Cas9.

| strain | transhemizygote parent | WT | transgenic | total | transmission rate ctrl (%) |
|---|---|---|---|---|---|
| Sco$^{GFP}$-CP | male | 451 | 588 | 1039 | 56.59 |
| | female | 382 | 427 | 809 | 52.78 |
| | overall | 833 | 1015 | 1848 | 54.92 |
| Aper1-Sco | male | 458 | 456 | 914 | 49.89 |
| | female | 515 | 491 | 1006 | 48.81 |
| | overall | 973 | 947 | 1920 | 49.32 |
| ScoG$^{CFP}$-AP2 | male | 268 | 245 | 513 | 47.76 |
| | female | 380 | 340 | 720 | 47.22 |
| | overall | 648 | 585 | 1233 | 47.45 |

**Table S5: Transmission rates and homing rates.**

| strain | transhemi-zygote parent | total progeny analyzed | excluded (Cas9+) | $E_{pos}$ | $E_{neg}$ | n | transmission rate (%) | homing rate (%) |
|---|---|---|---|---|---|---|---|---|
| Sco$^{GFP}$-CP | male | 265 | 118 | 143 | 4 | 147 | 97.28 | 94.56 |
| | female | 294 | 158 | 136 | 0 | 136 | 100.00 | 100.00 |
| | overall | 559 | 276 | 279 | 4 | 283 | 98.59 | 97.17 |
| Aper1$^{GFP}$-Sco | male | 315 | 126 | 182 | 9 | 191 | 95.29 | 90.58 |
| | female | 778 | 394 | 384 | 0 | 384 | 100.00 | 100.00 |
| | overall | 1029 | 520 | 566 | 9 | 575 | 98.43 | 96.87 |
| ScoG$^{CFP}$-AP2 | male | 1328 | n.a. | 1173 | 155 | 1328 | 88.33 | 76.66 |
| | female | 1236 | n.a. | 1158 | 78 | 1236 | 93.69 | 87.38 |
| | overall | 2564 | n.a. | 2331 | 233 | 2564 | 90.91 | 81.83 |
| ScoG-AP2 | male | 210 | n.a. | 208 | 2 | 210 | 99.05 | 98.10 |
| | female | 164 | n.a. | 164 | 0 | 164 | 100.00 | 100.00 |
| | overall | 374 | n.a. | 372 | 2 | 374 | 99.47 | 98.93 |

$E_{pos}$ and $E_{neg}$ refers to individuals with or without the effector construct, respectively. The homing rate e was calculated as follows: $e = (n*0.5–E_{neg})/(n*0.5)*100$.
